# Experimental adaptation of dengue virus 1 to *Aedes albopictus* mosquitoes by *in vivo* selection

**DOI:** 10.1101/2020.07.16.206524

**Authors:** Rachel Bellone, Sebastian Lequime, Henri Jupille, Giel P. Göertz, Fabien Aubry, Laurence Mousson, Géraldine Piorkowski, Pei-Shi Yen, Gaelle Gabiane, Marie Vazeille, Anavaj Sakuntabhai, Gorben P. Pijlman, Xavier de Lamballerie, Louis Lambrechts, Anna-Bella Failloux

## Abstract

In most of the world, Dengue virus (DENV) is mainly transmitted by the mosquito *Aedes aegypti* while in Europe, *Aedes albopictus* is responsible for human DENV cases since 2010. Identifying mutations that make DENV more competent for transmission by *Ae. albopictus* will help to predict emergence of epidemic strains. Ten serial passages *in vivo* in *Ae. albopictus* led to select DENV-1 strains with greater infectivity for this vector *in vivo* and in cultured mosquito cells. These changes were mediated by multiple adaptive mutations in the virus genome, including a mutation at position 10,418 in the DENV 3’UTR within an RNA stem-loop structure involved in subgenomic flavivirus RNA (sfRNA) production. Using reverse genetics, we showed that the 10,418 mutation alone does not confer a detectable increase in transmission efficiency *in vivo*. These results reveal the complex adaptive landscape of DENV transmission by mosquitoes and emphasize the role of epistasis in shaping evolutionary trajectories of DENV variants.

## Introduction

Vector-borne diseases represent almost one fourth of all emerging infectious diseases worldwide (1). Among the emerging diseases, arboviruses occupy the top stair with several million human cases reported annually (2). Dengue virus (DENV) belongs to the *Flavivirus* genus of the *Flaviviridae* family and is by far the most important arboviral disease, with the number of human dengue infection cases exceeding 300 million annually; 96 million are symptomatic dengue fever/hemorrhagic fever leading to an estimated 22,000 human deaths (3). DENV is comprised of four antigenically distinct but genetically related serotypes referred to as DENV1-4 (4). Clinical manifestations range from mild cases of dengue fever to severe cases of dengue hemorrhagic fever and/or dengue shock syndrome. All four DENV serotypes are now circulating in Asia, Africa and America (5). In past centuries, dengue was not an uncommon disease in Europe: the last record of a dengue outbreak in the 20^th^ century was in Athens, Greece, in 1927–1928 (6). This outbreak was unusual by the number of cases (~1 million) and the importance of severe clinical symptoms (e.g. hemorrhagic manifestations) leading to deaths (~1,000). After this Greek episode, dengue disappeared from Europe (7) as the mosquito *Aedes aegypti* disappeared from Eastern Mediterranean after 1935 through improving sanitation and mosquito control measures (8). No local transmission of DENV has been reported in Europe until 2010, when clusters of autochthonous cases were reported in Southern France (9) and Croatia (10). In France, several transmission episodes were successively reported: 2013-2015 (11–13), 2018-2019 (14, 15). The vector was *Aedes albopictus*, first detected in Europe in 1979 in Albania (16), then in 1990 in Italy (17), and today, established in more than 20 European countries (18). Unexpectedly, *Ae. albopictus* from France was shown to be more competent to experimentally transmit DENV-1 strains compared to its counterpart *Ae. aegypti* from the French West Indies (19). Contrary to *Ae. aegypti*, the mosquito *Ae. albopictus* which is native to South-East Asia, has a broader range of hosts (20). When *Ae. aegypti* is absent, *Ae. albopictus* can be responsible for DENV epidemics, as shown for the outbreaks in the Seychelles islands (21), Japan (22), La Réunion Island (23), and Hawaii (24). However, to date, *Ae. albopictus* is considered a minor vector of DENV relative to *Ae. aegypti* (25).

DENV serotypes have caused recent epidemics by changing their host ranges to increase infections in humans (26). As most arboviruses, DENV is capable of rapidly adapting to changes in their environment (or novel hosts) due to the accumulation of one or more specific mutations in the viral genome. For DENV, mutations in the 3’ untranslated region (UTR) have previously been linked to increased epidemiological fitness of the virus *via* a mechanism involving increased expression of 3’ UTR-derived subgenomic flavivirus RNA (sfRNA) expression (27). SfRNA is a known determinant for mosquito transmission of multiple flaviviruses like DENV, Zika virus and West Nile virus (28–32). Nucleotide substitutions in the 3’UTR reducing or ablating sfRNA expression negatively impact viral infection and transmission rates, suggesting that there is evolutionary pressure on conservation of RNA structures that dictate sfRNA expression in mosquitoes (33, 34). These studies indicate that subtle changes in the viral nucleotide composition can enhance the viral epidemic potential. On the same line, CHIKV has acquired the ability to spread globally owing to a single *Ae. albopictus*-adaptive mutation E1-226V (35). This mutation increased the infectivity of CHIKV in *Ae. albopictus* (36, 37).

We hypothesize that DENV can be selected for enhanced transmission by European *Ae. albopictus,* which would provide insight into future epidemic DENV strains that could pose a threat to human health. We conducted an experimental evolution study to identify nucleotide changes in the DENV genome by serially passaging DENV-1 isolates from Thailand (30A) and France (1806) in an *Ae. albopictus* population from Nice, France. Ten total passages were completed after which viral isolates were deep sequenced to identify newly acquired mutations. Importantly, we investigated whether the adaptation to the mosquito vector resulted in enhanced transmission potential or replication rate in mosquitoes. These results exemplify the potential of virus-adaptation studies for the identification of DENV strains likely to emerge.

## Results

### European *Ae. albopictus* are differentially susceptible to DENV-1

Arboviral transmission requires competent mosquitoes. To test whether European populations of *Ae. albopictus* can sustain local transmission of DENV-1, as reported in France (9) and Croatia (10), *Ae. albopictus* mosquitoes from Alessandria and Genoa (Italy), Cornelia and Martorell (Spain), Nice and Saint-Raphael (France) were experimentally infected with DENV-1 1806 from France or with DENV-1 30A from Thailand. Only engorged females were kept for analysis (samples size indicated in Fig. 1). When examining viral infection rate (Fig. 1a) and dissemination efficiency (Fig. 1b) at 14 and 21 days post-infection (dpi), percentages increased along with the dpi for the majority of virus/mosquito combinations. Within a mosquito population, the viral strain did not play a major role in either infection rate or dissemination efficiency (Fisher’s exact test: p > 0.05 after Bonferroni correction; Table S1). In contrast, we observed significant differences between mosquito populations (Fisher’s exact test: p < 0.05; Table S2) meaning that the geographic origin of the mosquito population is a critical factor that determines the outcome of viral infection and dissemination. Viral loads in heads (indicative of a successful dissemination from the midgut) did not mostly differ among mosquitoes having disseminated the virus (Fig. 1c, Table S3) (Wilcoxon Rank-Sum test: p > 0.05). No viral particles were detected in mosquito saliva (data not shown).

**Figure 1.**
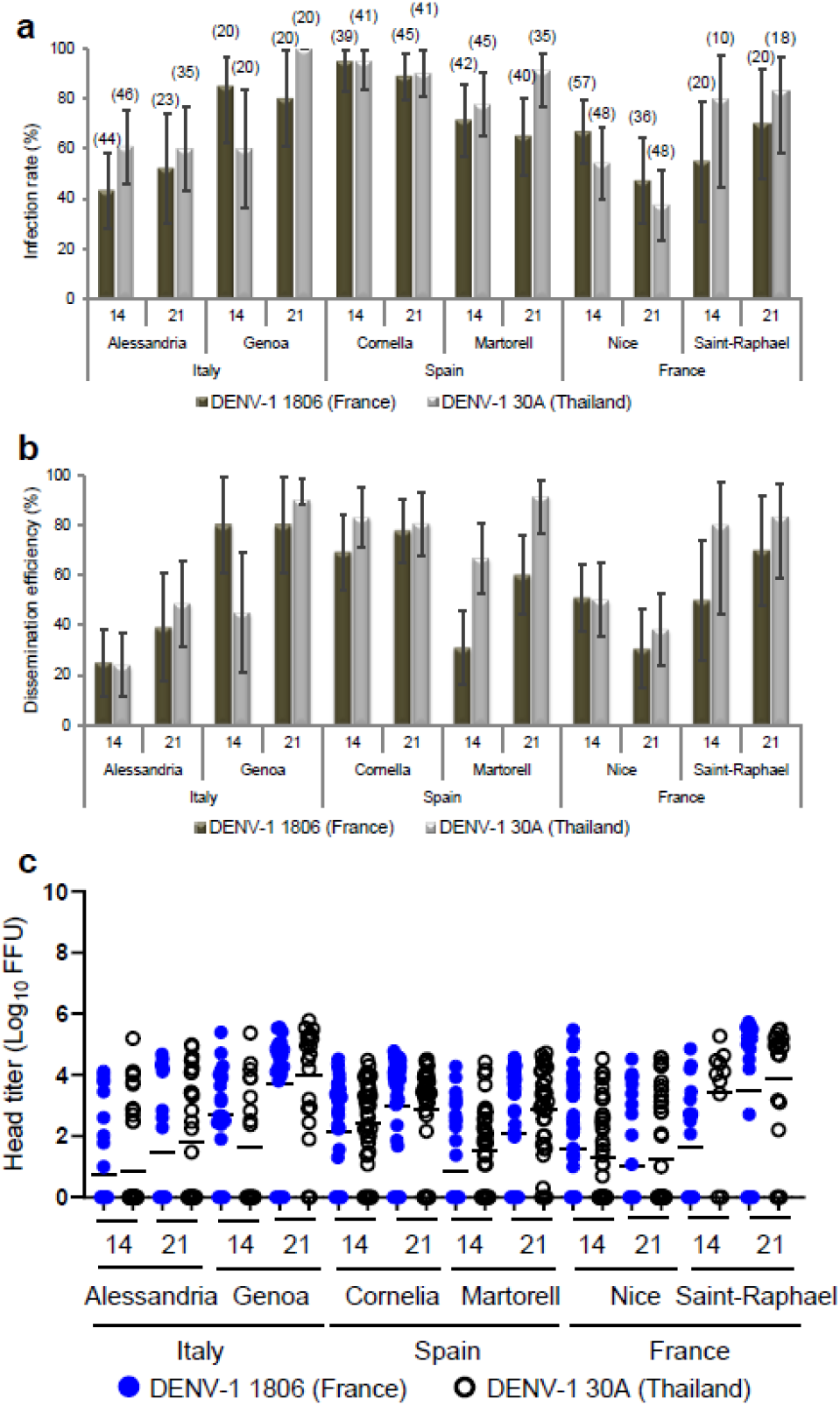
Susceptibilities of six European *Ae. albopictus* populations to DENV-1 (1806 and 30A): **a** infection rate, **b** dissemination efficiency and **c** viral titers in heads. Adult female mosquitoes were challenged with DENV-1 from France (1806) and Thailand (30A) at a titer of 10^7^ FFU/mL. At 14 and 21 dpi, mosquitoes were sacrificed and decapitated. Bodies and heads were homogenized and titrated on C6/36 cells. Infection rates were determined using positive/negative scoring (i.e. without estimating the number of viral particles), while viral titers at sites of dissemination were quantified *via* focus-forming assay. The error bars correspond to the confidence intervals (95%) (**a**, **b**), and the bar to the mean (**c**). In brackets is indicated the sample size.

### Experimental adaptation of DENV-1 to *Ae. albopictus*

To examine whether DENV-1 can adapt to *Ae. albopictus*, DENV-1 1806 and DENV-1 30A were passaged 10 times in duplicate in *Ae. albopictus* mosquitoes from Nice, France (Fig. 2). Viral titers of cell culture supernatants collected at each passage fluctuated slightly from passages 1 to 10 (Fig. S1). Additionally, the viruses were passaged 10 times in duplicate in *Ae. albopictus* C6/36 cells as a cell culture control. Full viral genomes were examined by deep sequencing at each passage (1–10) for the two replicates (R1 and R2) of *in vivo* mosquito infections and for passages 0 (parental strain), 1, 5 and 10 for the C6/36 cell culture control.

**Figure 2.**
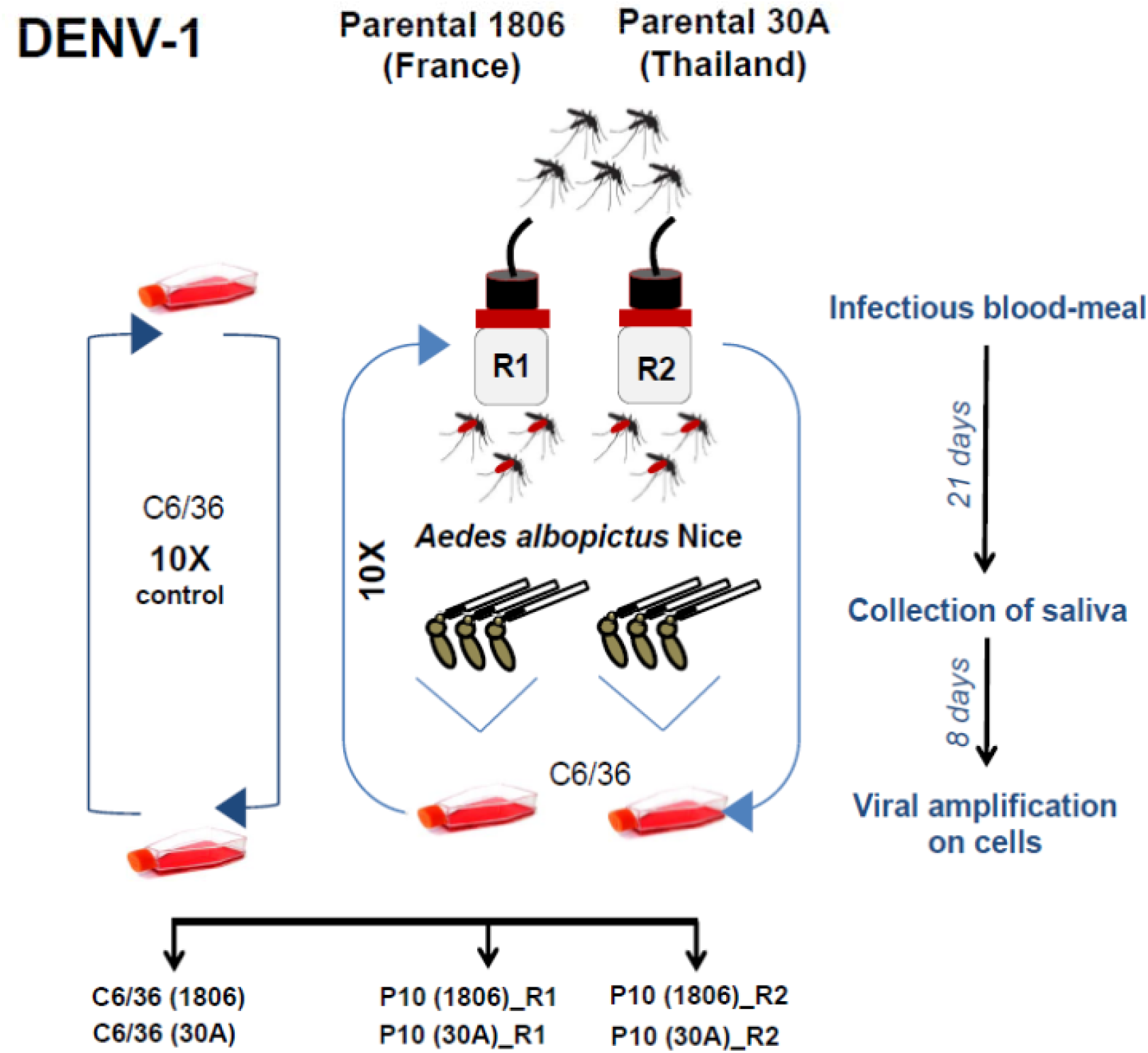
Experimental design for DENV-1 adaptation to *Ae. albopictus*. The parental strains 1806 (France) and 30A (Thailand) were passaged 10 times on a single *Ae. albopictus* population from Nice, France. Each passage includes: mosquito infectious blood-meal with DENV-1, collection of mosquito saliva at day 21 post-infection, viral amplification of saliva on *Ae. albopictus* cell cultures for 8 days, and initiation of the next passage using the viral suspension obtained. Control isolates were serially passaged 10 times on C6/36 cells. Two replicates R1 and R2 were performed.

The two parental DENV-1 strains, 1806 and 30A, yielded a mean sequencing depth of 68,687X (1806) and 133,941X (30A), covering 99.97% and 100% of the reference genome at >100X. With the exception of 30A (passage 4, R2 in mosquitoes; covering only 34.62% of the reference genome at >100X), all passages had a mean coverage between 995X and 211,830X, paving between 100 and 99.25% of the reference genome at >100X (Fig. S2).

No major changes in single nucleotide variants (SNV) frequencies were detected when DENV-1 isolate was serially passaged on C6/36 cells (Fig. 3a). Remarkably, we did not detect a single mutation that reached consensus level (frequency >50%) in the C6/36 control passages. In contrast, when DENV-1 1806 or 30A was passaged in *Ae. albopictus* mosquitoes, consensus level variants were detected as soon as passages 2 (DENV-1 1806) and 5 (DENV-1 30A), and a total of 30 consensus level variants were detected at passage 10 (Fig. 3a, Table S4). In total, twenty consensus level SNVs were detected in DENV-1 30A (positions 448, 694, 1611, 1768, 1959, 2002, 2200, 2716, 2977, 3442, 5822, 6658, 6728, 7267, 7952, 8149, 8485, 9504, 10208, 10258) and 10 in DENV-1 1806 (positions 1840, 2719, 3001, 3757, 4552, 4606, 5667, 7360, 9067, 10418). Out of the 30 consensus variants, 23 synonymous changes, 6 non-synonymous and one variant located in the 3’UTR were detected (Fig. 3b). The variant located at position 10,418 in the 3’UTR was the only SNV shared between the replicates R1 and R2 of DENV-1 1806. Its frequency increased over the passages, reaching consensus level at passage 4 for replicate 1 and passage 8 for replicate 2 (Fig. 3a). The variant became almost fixed (frequency > 99%) at passage 5 for replicate 1 and passage 10 in replicate 2. No SNV was common to the two replicates for DENV-1 30A. These results indicate that DENV-1 accumulates mutations during passaging in *Ae. albopictus* that likely facilitate virus replication in the mosquito or virus dissemination into the saliva to facilitate transmission.

**Figure 3.**
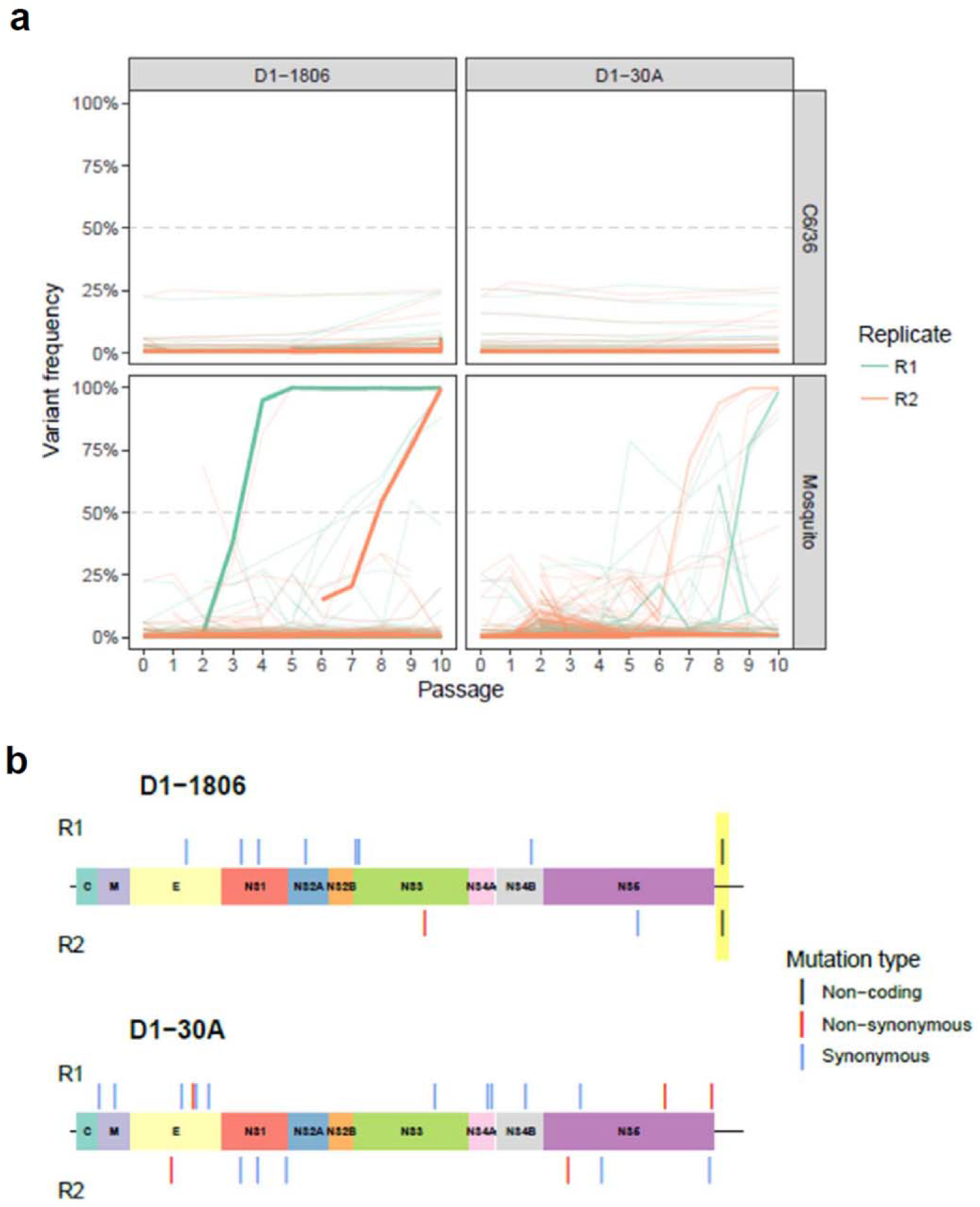
Frequency variation and genomic position of consensus level-reaching variants during passages. **a** The top panels correspond to DENV-1 passaged on C6/36 cells as controls and the bottom panels to DENV-1 passaged on *Ae. albopictus* mosquitoes. Bold lines represent the variant 10,418 in the 3’UTR. **b** Variants are represented with a colored segment according to the mutation type (non-coding: black; non-synonymous: red; synonymous: blue). The position of the only shared variant between two replicates is highlighted in yellow.

### *Ae. albopictus* adapted DENV-1 1806 has an increased transmission rate in *Ae. albopictus*

To investigate whether the mosquito adapted DENV-1 1806 has increased transmission potential as compared to the parental isolate, *Ae. aegypti* Pazar (Turkey) and *Ae. albopictus* Nice (France) were provided with an infectious blood-meal containing 10^7^ FFU/mL DENV-1 1806 parental, or replicate of the mosquito passaged virus (R1 and R2). Viral infection rates were high (>40%) and higher for the P10 viruses in *Ae. albopictus* at 21 dpi (Fig. 4a-b). Viral dissemination was lower at early dpi but remained high at 21 dpi in both mosquito species with a higher dissemination of P10 viruses compared to the parental virus (Fig. 4c-d). Transmission was surprisingly low in *Ae. aegypti* (Fig. 4E) compared to *Ae. albopictus* (Fig. 4f) suggesting a stronger effect of salivary glands as a barrier to virus release in saliva of *Ae. aegypti*. At 21 dpi, the two P10 viruses had higher infection rates (Fig. 4b), higher dissemination rates (Fig. 4d) and higher transmission rates (~2.5 fold) in *Ae. albopictus* than their parental strain (Fisher’s exact test: p < 10^−4^; IR, p = 0.0001; DE, p = 0.0001; TE, p = 0.022). Collectively, these results indicate that the accumulation of adaptive mutations during passaging in *Ae. albopictus* for the P10_R1 and P10_R2 viruses is beneficial for virus infection, dissemination and thus transmission by *Ae. albopictus* mosquitoes.

**Figure 4.**
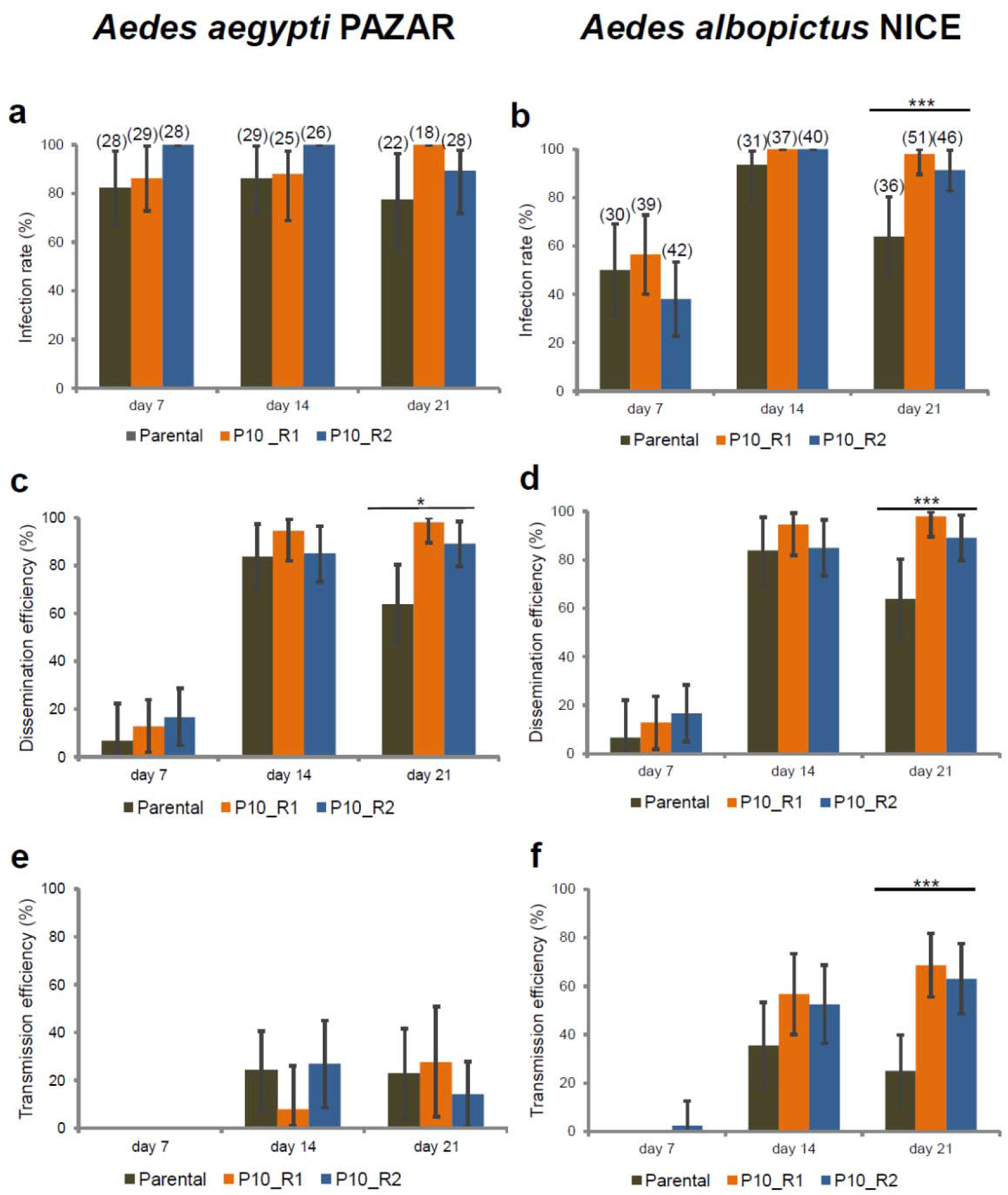
Infection, Dissemination and Transmission of DENV-1 (Parental, P10_R1, and P10_R2) by *Ae. aegypti* Pazar and *Ae. albopictus* Nice. Mosquitoes were exposed to blood meals at a titer of 10^7^ FFU/mL. Females were examined at 7, 14 and 21 dpi. Mosquito body (thorax and abdomen) and head were processed individually to determine (**a**, **b**) the infection rate (IR, proportion of mosquitoes with infected body among the engorged mosquitoes) and (**c**, **d**) the dissemination efficiency (DE, proportion of mosquitoes with infected head among tested mosquitoes). (**e**, **f**) Saliva was collected from individual females to determine the transmission efficiency (TE, proportion of mosquitoes with infectious saliva among tested mosquitoes). The Parental strain corresponds to DENV-1 1806, and P10_R1 and P10_R2 refer, respectively, to replicate 1 and replicate 2 of the 10^th^ *in viv*o passages of DENV-1 1806 on *Ae. albopictus*. Asterisks refer to a significant difference (***p < 10^−3^). In brackets, the number of mosquitoes tested. The error bars correspond to the confidence intervals (95%).

### *Ae. albopictus* adapted DENV-1 1806 has a replicative advantage in RNAi-competent and -deficient mosquito cells

To determine whether the adaptive mutations in DENV-1 1806 after serial passaging in *Ae. albopictus* mosquitoes were causing a replicative advantage, we examined the replication kinetics of DENV-1 1806 parental, R1 and R2 in *Ae. albopictus* C6/36 (RNAi-deficient) and U4.4 (RNAi-competent) cells (Fig. 5a-b) compared to kinetics in HFF cells (Fig. 5c). In C6/36 cells, the parental virus reached slightly lower titers at 6, 24 and 48 hours post-infection (hpi) as compared to the mosquito passaged R1/R2 viruses (Fig. 5a). The mosquito adapted R1/R2 viruses presented a significant increase in viral titer at 24 hpi (R1 (mean ± SD): Log_10_ 4.42 ± 0.10; R2: Log_10_ 4.68 ± 0.14) compared to the parental strain (Log_10_ 3.31 ± 0.15) (χ^2^ test: p = 0.027). In U4.4 cells, the same trend was observed (Fig. 5b) with a lower titer at 24 hpi for the parental strain (Log_10_ 2.84 ± 0.06) as opposed to the two P10 viruses (R1: Log_10_ 4.19 ± 0.29; R2: Log_10_ 4.47 ± 0.03). In human cells (Fig. 5c), viral titers remained between 2 and 3 Log_10_ from 0 to 72 hpi. These results indicate that the adaptive mutations after serial passaging of DENV-1 1806 in *Ae. albopictus* mosquitoes increase the replication rate in mosquito cells and a disadvantage in human cells.

**Figure 5.**
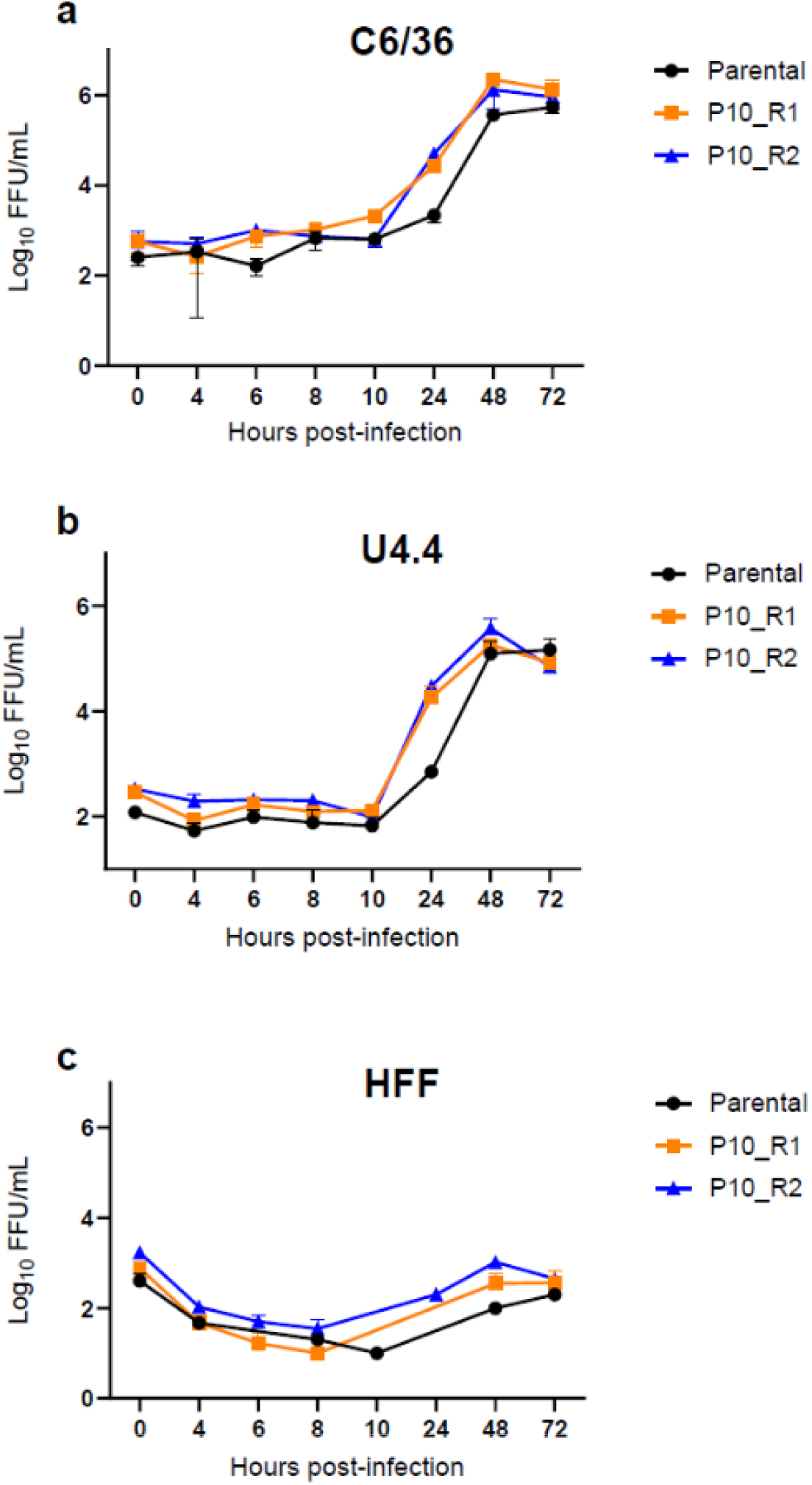
Growth curves of the two passages 10 of DENV-1 1806 strain in two cell lines, (**a**) *Ae. albopictus* C6/36 cells, (**b**) *Ae. albopictus* U4.4 cells, and (**c**) human foreskin fibroblasts HFF cells. Cells were infected with the parental strain and the two replicates of the 10^th^ passages of DENV-1 1806 (P10_R1 and P10_R2) at a MOI of 0.1. Supernatants were collected at 4, 6, 8, 10, 24, 48 and 72 hrs post-inoculation. The number of infectious viral particles was determined by focus fluorescent assay on *Ae. albopictus* C6/36 cells. Three replicates were performed for each cell-virus pairing. Error bars show standard deviations.

### Vizualisation of substitutions in 3’UTR of DENV-1 1806 on RNA stem-loop structures

The highly structured flavivirus 3’UTR is important for virus replication, genome translation and production of non-coding sfRNA (38). sfRNA is formed as a result of incomplete degradation of the viral genomic RNA by the 5’-3’ exoribonuclease XRN1, which stalls on stem loop (SL) and dumbbell (DB) RNA structures in the 3’UTR (38, 39). It has been shown that passaging DENV on mosquito cells can result in high mutation rates in the 3’UTR and might alter the abundance of sfRNA during infection (33, 40). The largest sfRNA species, sfRNA1, determines pathogenicity (41), inhibits host innate immunity (27, 42) and is essential for efficient transmission of flaviviruses by mosquitoes (28–32, 43, 44). We therefore investigated if the consensus level mutation 10,418 that occurred in the 3’UTR after passaging could lead to changes in the 3’UTR secondary RNA structures and subsequent sfRNA formation. Mutations with an SNV frequency ≥0.05 only occurred in the SL-II and 3’SL structures (Fig. 6a; red nucleotides). When examining mutations in the 3’SL, the same SNVs were found in the two parental strains and the passages 10 indicating that those mutations were already present in the initial viral populations and were not selected consequently to serial passages in mosquitoes (Fig. 6b). We observed that the mosquito passaged DENV-1 1806 presented a U→C substitution at position 10,418 on the top of SL-II, which was not observed for DENV-1 30A. SL-II is the XRN1 stalling structure required for sfRNA2 formation, which requires the presence of a RNA pseudoknot interaction and a complex tertiary folding (45) (Fig. 6b). Pseudoknot formation and other known tertiary RNA interactions are not expected to be directly disrupted due to the 10,4018 sequence change (Fig. 6b). However, the U→C mutation may indirectly affect the 3D folding of the stem loop. Passages on C6/36 cells also gave rise to lower-frequency mutations in SL-II (frequency < 0.2), but none of them reached consensus level.

**Figure 6.**
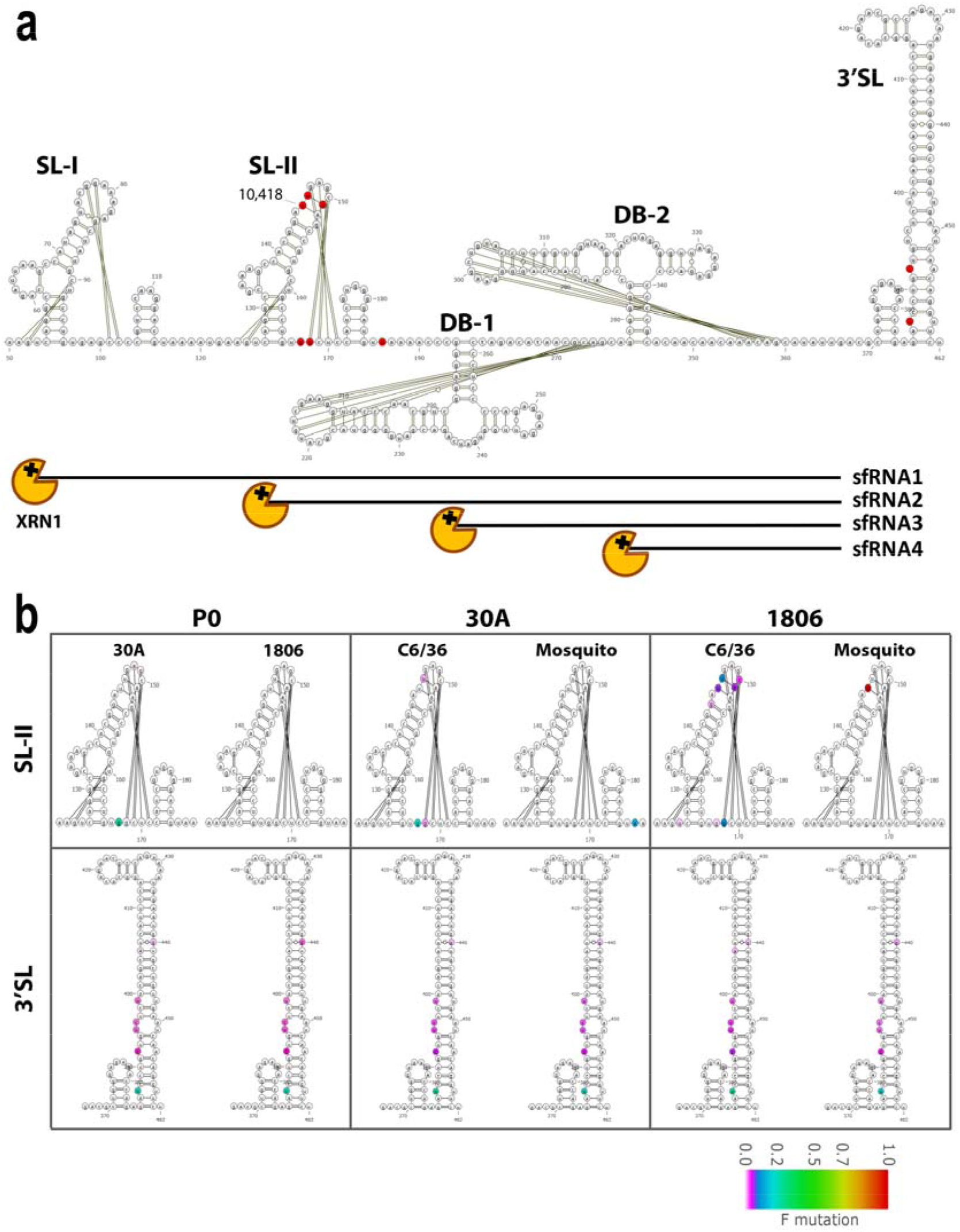
Analysis of mutation frequencies in the DENV-1 3’UTR. **a** Schematic overview of the DENV-1 3’UTR secondary RNA structure, indicating from 5’ to 3’ the stem loop (SL)-I, SL-II, dumbbell (DB)-1, DB-2 and 3’SL RNA structures. Single nucleotide variants with a frequency ≥0.05 after 10 passages of DENV-1 1806 or 30A in either C6/36 or mosquitoes are highlighted in red. Pseudoknots and other tertiary RNA interactions are indicated by the black lines. **b** Analysis of the mutation frequencies in SL-II and the 3’SL of the parental DENV-1 30A and 1806 sequences, and passages P10 (1806) and P10 (30A). The mutation frequency is indicated by color on a scale from 0 (white) to 1 (red).

### Increased transmission potential of *Ae. albopictus*-adapted DENV-1 is not associated with significantly increased sfRNA production

As sfRNAs are generated due to stalling of XRN1 on RNA secondary structures in the viral 3’UTR (41), sequence changes in the 3’UTR, in particular those that occur in RNA structures involved in XRN1 stalling, may affect the length and expression level of sfRNAs. To investigate whether the production of sfRNA is affected by the mutation 10,418 in DENV-1 1806 R1/R2, a Northern blot analysis was performed using a 3’UTR specific probe on total RNA extracted from U4.4 cells infected with DENV-1 1806 parental, R1 or R2 (Fig. 7). Viral gRNA and abundant sfRNA1 were produced by both the parental and R1/R2 viruses (Fig. 7a). The quantity of sfRNA1 was visually similar across all samples on the gel, although ImageJ quantification of the band intensities revealed that the ratio of sfRNA/gRNA was ~1.5 fold higher in R1 and R2 samples as compared to the parental samples (Fig. 7b). Minimal amounts of smaller sfRNA species (i.e. sfRNA2, 3, 4) were observed, indicating that DENV-1 predominantly produces sfRNA1 during infection of mosquito cells. These results show that the increased transmission potential of the *Ae. albopictus* adapted DENV-1 1806 is unlikely to be caused by differences in sfRNA production.

**Figure 7.**
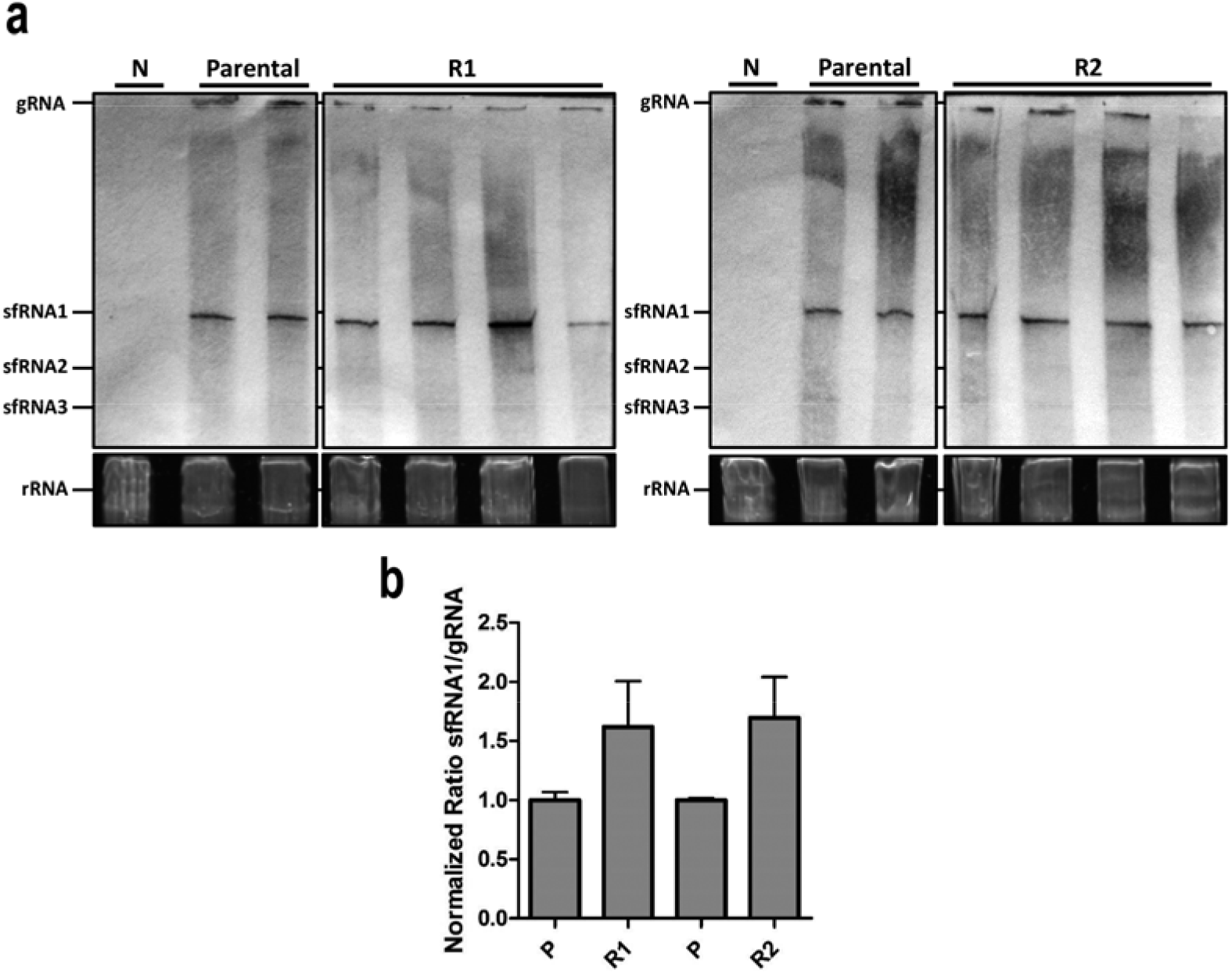
Northern blot detection of sfRNA production after C6/36 cells infection with DENV-1 1806. **a** Visualization of sfRNA. 5μg of total RNA from the parental and the two replicates P10_R1 and P10_R2 of DENV-1 1806 or non-infected cells (N) was size separated on a 6% Polyacrylamide/Urea gel. One gel was run for the R1 samples (left) and one for the R2 samples (right), including the negative and parental samples on each gel. Then, RNAs were blotted onto Hybond-N paper and subjected to northern-blotting with a DENV-1 3’UTR specific probe. The bands shown correspond to the DENV genomic RNA (gRNA) and subgenomic flavivirus RNA (sfRNA). As loading control, the ribosomal RNA (rRNA) from the EtBr stained gel are shown. **b** Quantification of the ratio of sfRNA to gRNA production. Band intensities were determined using the ‘Measure’ function in ImageJ. The intensity of the background (lane N) was subtracted from the readings before the ratio sfRNA/gRNA was calculated by dividing the intensity of the sfRNA by the intensity of the gRNA band for each sample, and then normalized to the average ratio of the parental samples. The statistics were performed using a two-tailed unpaired t-test.

### The 10,418 mutation alone does not significantly enhance DENV-1 transmission by *Ae. albopictus*

To test whether the U→C substitution at position 10,418 alone would recapitulate the observed phenotype in *Ae. albopictus*, three reverse genetic constructs (Parental construct, P10 construct 1 and P10 construct 2) were produced using the ISA method and then sequenced. As expected, the two P10 constructs presented the 10,418 mutation at a frequency close to 100% (Table S5). The genetic constructs also displayed other SNVs at frequencies higher than 5% but none of them reached consensus level with the exception of one SNV with 52.5% frequency in P10 construct 1 (5173/nsp3) and two SNVs close to fixation in P10 construct 2 (7321/nsp4b and 9571/nsp5).

Twenty-one days after an infectious blood meal containing the reverse genetic constructs, *Ae. albopictus* Nice mosquitoes were examined for transmission by analyzing viral particles in their saliva. When estimating the transmission efficiency, no significant differences were detected when comparing the five viral strains, the reverse genetic constructs in reference to the template (Parental, P10) and the two P10 constructs (1 and 2) (Fisher’s exact test: p > 0.05) (Fig. 8a). Similarly, when examining the number of viral particles in individual mosquito saliva, no statistical significance was found whatever the comparison (Wilcoxon Rank-Sum test: p > 0.05) (Fig. 8b).

**Figure 8.**
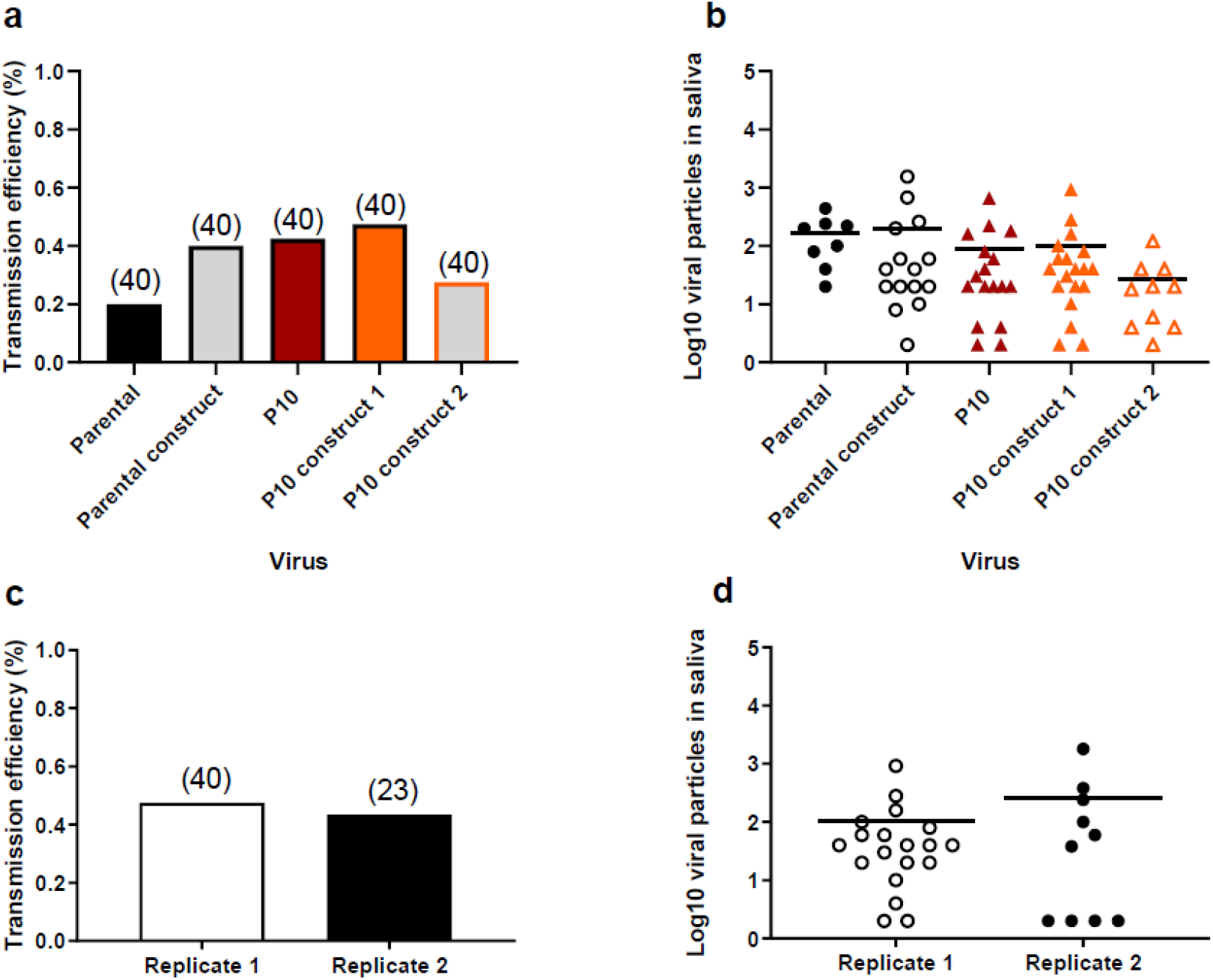
Reverse genetic constructs with the 10,418 mutation do not show higher transmission in *Ae. albopictus*. **a** Transmission of reverse genetic constructs (Parental construct, P10 construct 1, P10 construct 2) by *Ae. albopictus* Nice with reference to Parental and P10 strains. Twenty-one days after an infectious blood meal at a titer of 10^7^ FFU/mL, mosquitoes were processed for saliva collection to determine the transmission efficiency (TE, proportion of mosquitoes with infectious saliva among tested mosquitoes). **b** Viral loads in saliva were estimated by focus fluorescent assay on *Ae. albopictus* C6/36 cells. **c, d** A second replicate using the P10 construct 1 was performed. Bars indicate the mean.

To confirm the profile of P10 construct 1, we performed a replicate using the same experimental design. TE (Fig. 8c) and the viral load in saliva (Fig. 8d) were determined. We found that the replicate 2 shares the same profile than the replicate 1 (Fig. 8a, 8b). Altogether, these results indicate that the mutation 10,418 alone does not enhance transmission of DENV-1 in *Ae. albopictus*.

## Discussion

In this study, we developed an experimental evolution approach to enhance the transmission of DENV-1 by the vector *Ae. albopictus*. A number of conserved nucleotide variants were observed in all mosquito-passaged DENV-1 viruses, but not in the viruses passaged in mosquito cells, indicating that DENV-1 virus adapts specifically to cope with the adaptive pressure in the mosquito. A nucleotide change at position 10,418 in the 3’UTR was observed in both replicates of the *Ae. albopictus* adapted DENV-1 1806. Independent fixation of the mutation in two replicates with a sharp rise in frequency is consistent with adaptive evolution. Importantly, the *Ae. albopictus* adapted isolates displayed higher DENV-1 infection, dissemination and transmission efficiencies in *Ae. albopictus*. Our results show that we have successfully adapted DENV-1 to *Ae. albopictus*, through selection of adaptive mutations including the 10,418 mutation in the 3’UTR of the viral genome by sequential passaging *in vivo*.

*Ae. albopictus* usually acts as a secondary vector of DENV (20), but in the absence of *Ae. aegypti*, it can act as the main vector in some regions including Europe (9–13). First detected in Albania in 1979 (16), *Ae. albopictus* is now present in more than 20 European countries (18). We showed that *Ae. albopictus* from France, Italy and Spain were susceptible to infection by DENV-1 (Fig. 1), indicating that *Ae. albopictus* is indeed a vector species for DENV in Europe. The main sources of introductions in Europe were mosquitoes from Italy, which were previously imported from North America (46). Recurrent introduction events have contributed to increase the genetic diversity of European *Ae. albopictus* populations (47), an important factor shaping vector competence (25).

Here, we experimentally selected DENV-1 isolates for enhanced transmission by *Ae. albopictus*. Our experimental procedure was designed to accelerate the selection process of DENV-1 by serial passages in *Ae. albopictus* mosquitoes without alternation in the mammalian host. After 10 passages in *Ae. albopictus* collected in Nice, France, we successfully adapted DENV-1 1806 and DENV-1 30A to *Ae. albopictus* through the accumulation of adaptive mutations across the genome, although only a single mutation was fixed in both replicates for the 1806 isolate. Importantly, for DENV-1 1806, these adaptive mutations increased the infection, dissemination and transmission rates of DENV-1 by *Ae. albopictus* (Fig. 4). Furthermore, growth kinetics of the DENV-1 1806 viruses were increased in both RNAi-competent U4.4 and RNAi-deficient C6/36 cells, indicating that the mutations cause an increase in viral replicative fitness in cell cultures regardless of a functional RNAi machinery (Fig. 5). These mosquito-selected viral variants were less adapted to replicate on mammalian cells (48). In a similar approach, Stapleford et al. (2014) succeeded in monitoring the selection of epidemic variants of CHIKV adapted to *Ae. albopictus* consolidating the idea that *in vivo* approaches can contribute in predicting new variants able to emerge and displace currently circulating viral strains (49).

We identified a mutation that was fixed in both replicates of the mosquito passaged DENV-1806 isolate, suggesting that this residue is involved in the adaptive-evolution that results in the increased transmission and replication potential of the virus. This shared substitution was located at position 10,418 in the highly structured 3’UTR of the DENV-1 genome. Specifically, the mutation was present in the SL-II RNA structure (Fig. 6); which is required for the production of sfRNA2 (41). XRN1 stalls at SL and dumbbell (DB) RNA structures within the 3’UTR, which results in accumulating sfRNAs of different sizes (41, 50). The stalling of XRN1 occurs due to steric hindrance caused by interactions of pseudoknots (PK) and other tertiary RNA structures (45, 51). Prediction of RNA structures involved in XRN1 stalling (the so-called xrRNAs) with Mfold has proven to be an useful starting point but undeniably has limitations, e.g. pseudoknots cannot be predicted and 3D RNA folding is not taken into account. Although Mfold predictions and visual pseudoknot mapping have helped to elucidate mechanisms of XRN1 stalling (41, 45, 50), the exact structural basis for XRN1 stalling, the involvement of a unique three-way junction and internal tertiary interactions were only revealed by determining the crystal structure of several xrRNAs (52, 53). In mammalian cells, sfRNA is essential for inducing pathogenicity (41), and acts as an antagonist of innate immune responses (27, 42). In mosquito cells, sfRNA has been reported as an antagonist of the RNAi response *in vitro* (54, 55) and contributes to enhance the *in vivo* infection of mosquitoes and further dissemination from the midgut into the haemocoel (32) and subsequent salivary gland infection (44). Villordo et al. (2015) previously demonstrated that when passaging DENV-2 20 times in C6/36 mosquito cells, SL-II is highly mutated while the upstream SL-I mutates mostly upon passaging in mammalian cells (40). The mutations in SL-II were shown to increase DENV-2 replication in mosquito cells (40). The mutation that we found at the position 10,418 in SL-II is in line with these findings, supporting the mutation pressure on SL-II *in vivo*, although we did not observe significant mutations during passaging in C6/36 cells.

For DENV-2, it has been shown that during replication in human cells, mainly sfRNA1 is produced, while mosquito-adapted DENV-2 accumulates more abundant sfRNA3 and sfRNA4 (33). We show that DENV-1 1806 produced abundant sfRNA1 while quantities of sfRNA2,3 and possibly sfRNA4 were below the detection limit (Fig. 7), suggesting possible differences in the production of sfRNA species between DENV-1 and DENV-2. Despite the presence of the 10,418 mutation, the *Ae. albopictus* adapted DENV-1 1806 did not show a significantly altered production of sfRNA species. Although we cannot exclude an effect on cellular binding partners that might require an intact 3’UTR for their interaction with the viral genome (38), it is unlikely that sfRNAs were a primary driver of DENV-1 adaptation to *Ae. albopictus*.

We used reverse genetics to evaluate the effect of the 10,418 mutation on DENV-1 transmission by *Ae. albopictus* mosquitoes *in vivo*, but our results did not provide experimental support for a phenotypic effect of the 10,418 mutation alone. For two different genetic constructs harboring the 10,418 mutation (together with different adventitious mutations), there was no detectable difference in transmission efficiency. Introducing the 10,418 did not recapitulate the adapted phenotype of the P10 viruses and points to a more complex adaptive landscape than a single-mutation effect. Because our genetic constructs focused on the 10,418 mutation did not include other mutations present in the P10 viruses, it implies that the enhanced transmission phenotype reflected the combined effect of several mutations. Such epistatic relationships have been documented to shape the adaptive landscapes of CHIKV (56, 57) and more recently DENV (58). Interestingly, the 10,418 mutation was the only shared mutation among replicates of adapted viruses (Fig. 3), indicating that DENV-1 adaptation to *Ae. albopictus* can result from distinct evolutionary trajectories involving different sets of mutations.

Our experimental approach has succeeded in enhancing the transmission of DENV-1 by multiple passages in the *Ae. albopictus* vector. This protocol can be extended to other arboviruses and vector species, and contribute to predict future epidemic variants. Together, this may ultimately lead to new insights into the mechanisms of arbovirus transmission by mosquitoes.

## Materials and Methods

### Ethics statement

The Institut Pasteur animal facility has received accreditation from the French Ministry of Agriculture to perform experiments on live animals in compliance with the French and European regulations on care and protection of laboratory animals (EC Directive 2010/63, French Law 2013-118, February 6th, 2013). This study was approved by the Ethics Committee #89 and registered under the reference APAFIS#6573-201606l412077987 v2. Mice were only used for mosquito rearing as a blood source, according to approved protocol.

### Cell cultures

*Ae. albopictus* C6/36 cells were maintained at 28°C in Leibovitz L-15 medium supplemented with non-essential amino-acids (NEAA) (1X), 10% fetal bovine serum (FBS), 100 units/mL penicillin and 100 μg/mL streptomycin. These cells are defective in typical siRNAs, the hallmark of exogenous RNAi mediated antiviral immunity (59). *Ae. albopictus* U4.4 cells were maintained in L-15 medium supplemented with non-essential amino-acids (1X), 10% FBS, 100 units/mL penicillin and 100 μg/mL streptomycin at 28°C. HFF (Human Foreskin Fibroblast) cells were maintained at 37°C, 5% CO_2_ in Dulbecco’s Modified Eagle medium (DMEM) supplemented with pyruvate, 10% FBS, 100 units/mL penicillin and 100 μg/mL streptomycin. The human embryonic kidney HEK-293 cells (ATCC number CCL-1573) were grown at 37°C with 5% CO2 in tissue-culture flasks with vented caps, in a minimal essential medium (MEM, Life Technologies) supplemented with 7% FBS, 1% PS and 1X NEAA.

### Viruses

We used two DENV-1 strains isolated from DF cases: DENV-1 1806 from an autochthonous case from Nice, France in 2010 (provided by the National Reference Center of Arboviruses, France) and DENV-1 30A from a patient in Kamphaeng Phet, Thailand in 2010 (provided by the Afrims, Thailand and under accession number HG316482 in GenBank). The 2^nd^ passage of DENV-1 1806 on African green monkey kidney Vero cells (60) and the 2^nd^ passage of DENV-1 30A on C6/36 *Ae. albopictus* cells (61) were used for mosquito infections. Serial dilutions were used to determine the titer of viral stocks that was expressed in focus-forming units (FFU)/mL.

### Mosquito strains

Six populations of *Ae. albopictus* have been established from eggs: Genoa (Italy), Alessandria (Italy), Cornella (Spain), Martorell (Spain), Nice Jean Archet (France), and Saint-Raphael (France) (Table 1). They were tested to appraise vector competence to DENV-1 isolates. Together with *Ae. albopictus* Nice Jean Archet (France), *Ae. aegypti* Pazar (Turkey) was utilized to compare vector competence using viruses isolated after 10 passages on *Ae. albopictus*. Eggs were collected from ovitraps and sent to the Institut Pasteur in Paris, where they were reared in standardized conditions. After hatching, larvae were distributed in pans containing a yeast tablet renewed as needed in 1 L of tap water. Adults were placed in cages maintained at 28±1°C, at relative humidity of 80% and a light:dark cycle of 16h:8h, with free access to 10% sucrose solution. Oral infection experiments were performed using mosquitoes from the F2-F11 generations. Owing to the limited number of mosquitoes, only one biological replicate was performed for each pairing population-virus.

**Table 1.**
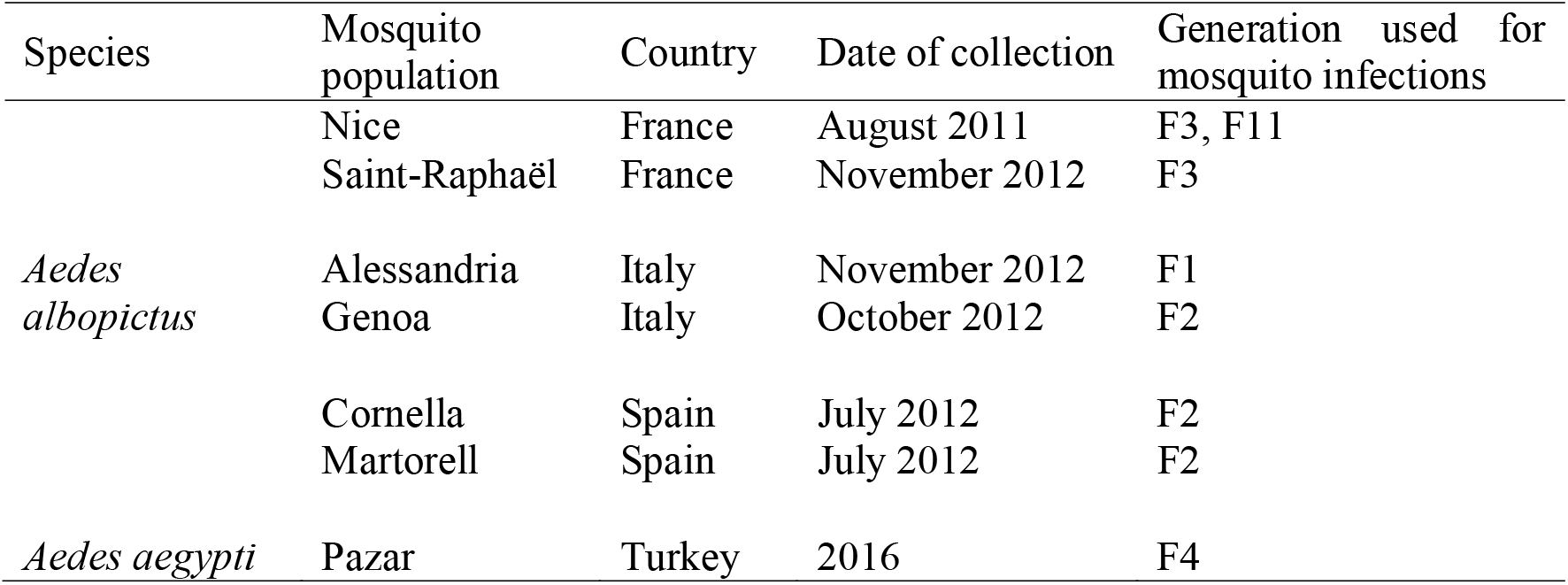
Details on mosquito populations used for experimental infections with DENV-1.

### Mosquito infections

One-week-old females were starved 24 hrs prior an infectious blood-meal in a BSL-3 laboratory. Five batches of 60 mosquito females were then allowed to feed for 15 min through a piece of pork intestine covering the base of a Hemotek feeder containing the infectious blood-meal maintained at 37°C. Only engorged females were kept and incubated under controlled conditions (28±1°C, relative humidity of 80%, light:dark cycle of 16h:8h).

#### For vector competence assays

Fourteen and 21 days after an infectious blood-meal provided at a titer of 10^7^ FFU/mL, vector competence was assessed based on two phenotypes: (i) viral infection of mosquito and (ii) viral dissemination from the midgut into mosquito general cavity. Infection rate (IR) was determined as the proportion of mosquitoes with infected midgut and dissemination efficiency (DE) was defined as the percentage of mosquitoes with virus detected in heads suggesting a successful viral dissemination from the midgut. IR and DE were calculated by titrating body and head homogenates.

#### For serial passages

As the first autochthonous DENV cases were reported in Nice in 2010 (9), *Ae. albopictus* isolated in Nice was used to achieve the experimental selection of DENV-1 isolates (Fig. 2). Mosquitoes were orally infected with DENV-1 supernatant provided in a blood-meal at a final titer of 10^6.5^ FFU/mL using the hemotek system. Engorged mosquitoes were incubated at 28°C for 19-21 days and then processed for saliva collection. 15-25 saliva were pooled and the volume of the pool was adjusted to 600 μL with DMEM prior to filtration through a Millipore H membrane (0.22 μm). An aliquot of 300 μL of each sample was used to inoculate a sub-confluent flask (25 cm^2^) of C6/36 *Ae. albopictus* cells. After 1 hr, the inoculum was discarded and cells were rinsed once with medium. Five mL of DMEM medium complemented with 2% FBS was added and cells were incubated for 8 days at 28°C. Cell culture supernatants were then collected and provided to mosquitoes to run the next passage. Passages P1 to P3 were performed with mosquitoes of the F3 generation and passages P4 to P10 with mosquitoes of the F4 generation. C6/36 supernatants collected at each passage were used undiluted for the next mosquito blood-meal. Ten passages were performed. Control isolates corresponded to serially passaged viruses on C6/36 cells to identify mutations resulting from genetic drift or adaptation to insect cell line; 500 μL of the previous passage were used to inoculate the next flask of C6/36 cells. Two biological replicates R1 and R2 were performed to test the variability between samples submitted to the same protocol of selection. Vector competence using the parental and P10 isolates was assessed by calculating: (i) infection rate (IR, proportion of mosquitoes with infected midgut), (ii) dissemination efficiency (DE, proportion of mosquitoes able to disseminate the virus from the midgut among tested mosquitoes), and (iii) transmission efficiency (TE, proportion of mosquitoes with the virus detected in saliva among tested mosquitoes).

### Virus deep sequencing

Total RNA was extracted from cell culture supernatant using QIAamp Viral RNA Mini Kit (Qiagen, Germany) and DNAse treated (Turbo DNAse, Life Technologies, USA). Following purification with magnetic beads (Agencourt RNAClean XP, Beckman Coulter, California, USA), RNA was reverse transcribed using Transcriptor High Fidelity cDNA Synthesis Kit and a specific 3’-UTR DENV-1 primer (Roche Applied Science, Mannheim, Germany), d1a5B 5’-AGAACCTGTTGATTCAACRGC-3’ (62). Second strand was then synthetized in a unique reaction with *E. coli* DNA ligase (New England Biolabs, Massachusetts, USA), *E. coli* DNA polymerase I (New England Biolabs), *E. coli* RNAse H (New England Biolabs) in second strand synthesis buffer (New England Biolabs). After purification with magnetic beads (Agencourt AMPure XP, Beckman Coulter), dsDNA was quantified with fluorometric method (Quant-iT PicoGreen dsDNA, Invitrogen, Massachusetts, USA).

Sequencing libraries were prepared using Nextera XT DNA Library Preparation Kit (Illumina, San Diego, USA), multiplexed and sequenced in single end in two independent runs on an Illumina NextSeq 500 platform using a mid-output 150-cycle v2 kit (Illumina). Reads were trimmed (Trimmomatic v0.33) (63) after demultiplexing (bcl2fastq v.2.15.0, Illumina) to remove adaptor sequences, and reads shorter than 32 nucleotides were discarded. Full-length genome of the DENV-1 1806 was assembled *de novo* using Ray v2.0.0 (64) with the original stock sample. The newly assembled DENV genome contig was extended in 3’ and 5’ using closest BLAST hit full DENV-1 genome (accession number EU482591). This chimeric construct was used to map reads used for assembly using Bowtie 2 v2.1.0 (65). Alignment file was converted, sorted and indexed using Samtools v0.1.19 (66). Coverage and sequencing depth were assessed using bedtools v2.17.0 (67). Single nucleotide variants and their frequency were called using LoFreq* v2.1.1 (68) and used to correct the chimeric construct. Only nucleotides with >10X coverage were conserved for generating the consensus sequence. A final full-length genome sequence for DENV-1 1806 strain was deposited to GenBank (accession number MG518567).

After quality control, reads from all samples were mapped to the newly assembled DENV-1 1806 strain genome sequence or previously sequenced reference genome KDH0030A (accession number HG316482) using Bowtie v2.1.0 (65). The alignment file was converted, sorted and indexed using Samtools v0.1.19 (66), and the coverage and sequencing depth were assessed for each sample using bedtools v2.17.0 (67). Single nucleotide variants (SNVs) and their frequency were then called using LoFreq* v2.1.1 (68), and their effect at the amino-acid level was assessed by SNPgenie v1.2 (69).

### RNA structure modeling in silico

The Mfold Web server was used with standard settings and flat exterior loop type (70) to fold the secondary RNA structures, which were then visualized using the VARNA RNA editing package (71). Pseudoknot RNA interactions were drawn as previously described for DENV (45, 72). Mutation frequencies of individual nucleotides were determined by averaging the nucleotide allele frequency from the deep sequencing results of the duplicates per treatment.

### Virus growth curves

To measure viral replicative fitness, growth curves were conducted in *Ae. albopictus* C6/36 and U4.4 mosquito cells, and Human Foreskin Fibroblasts (HFF) cells. Confluent cell monolayers were prepared and inoculated with viruses simultaneously in triplicates at a MOI of 0.1 PFU/cell. Cells were incubated for 1 hr in appropriate conditions and viral inoculum was removed to eliminate free virus. Five mL of medium supplemented with 2% FBS were then added and mosquito cells were incubated at 28°C (mosquito cells) or 37°C (human cells). At various times (4, 6, 8, 10, 24, 48 and 72 hrs) post-inoculation (pi), supernatants were collected and titrated by focus fluorescent assay on *Ae. albopictus* C6/36 cells. After incubation at 28°C for 5 days, plates were stained using hyper immune ascetic fluid specific to DENV as primary antibody (Millipore, Molsheim, France). A Fluorescein-conjugated goat anti-mouse was used as the second antibody (Thermofisher). Three viral strains were used: the parental strain and two 10^th^ passages, P10_R1 and P10_R2. Viral titer was expressed in FFU/mL. Three biological replicates were performed for each cell-virus pairing.

### RNA isolation and Northern blotting

Total RNA was isolated from cell monolayers using TRIzol reagent (Invitrogen, Massachusetts, France) following the manufacturer’s protocol. Mosquito DENV-1 infected bodies were homogenized individually in 500 μL of Leibovitz L15 medium (Invitrogen) supplemented with 2% fetal bovine serum for 1 min at maximum speed. Homogenates were then filtered with a filter unit (0.22 μm) (Ultrafree® MC-GV, Merck, New Jersey, USA). Two samples of each filtrate were inoculated onto monolayers of *Ae. albopictus* C6/36 cell culture in 6-well plates. After incubation at 28°C for 6 days, samples were homogenized with 1 mL TRIzol reagent. RNA isolations were performed using the standard TRIzol protocol. Samples were eluted in 30 μL RNase-free Milli-Q water and stored at −80°C until further processing. A DENV-1 3’UTR specific probe was generated by PCR reaction with GoTaq Polymerase (Promega, Wisconsin, USA) containing DIG DNA-labelling mix (Roche) and primers DENV-1 3’UTR FW (AGTCAGGCCAGATTAAGCCATAGTACGG) and DENV-1 3’UTR RV (ATTCCATTTTCTGGCGTTCTGTGCCTGG) using cDNA from cells infected with DENV-1 1806 as a template. Five micrograms of total RNA was subjected to sfRNA-optimized northern blot as has been described previously (32). Briefly, total RNA was denatured and size separated on 6% polyacrylamide–7 M urea–0.5× Tris-borate-EDTA (TBE) gel for 1.45 hrs at 150 V. The RNA was semi-dry-blotted on a Hybond-N membrane, UV cross-linked and pre-hybridized for 1 hr at 50°C in modified Church buffer containing 10% formamide. DENV-1 3’UTR specific Dig-labelled probe was denatured and blots were hybridized overnight at 50°C in modified church/10% formamide buffer containing 2 μL of DIG-labelled probe. Blots were developed with AP-labeled anti-DIG antibodies and NBT-BCIP solution before observing the signal using a Bio-Rad Gel Doc scanner. Quantification of band signal intensities was performed in ImageJ by transforming the image to 8-bit format, inverting the image, and analyzing the band intensity using the measure function. The Ratio sfRNA/gRNA was calculated by dividing the intensity of the sfRNA by the intensity of the gRNA band for each sample, and then normalized to the average ratio of the parental samples.

### ISA reverse genetics

The T>C mutation at position 10,418 identified at passage 10 was inserted into a DENV-1 1806 backbone using the ISA (Infectious Subgenomic Amplicons) reverse genetics method as previously described (73).

#### Preparation of subgenomic DNA fragments

The viral genome was amplified by RT-PCR from the DENV-1 1806 viral RNA as three overlapping DNA fragments. Two additional fragments were *de novo* synthesized (Genscript) and amplified by PCR (primers are listed in Table S6). The first primer consisted of the human cytomegalovirus promoter (pCMV) and the second primer of the last 367 nucleotides of the 3’UTR of the DENV-1 1806 with or without the 10,418 T>C mutation and the hepatitis delta ribozyme followed by the simian virus 40 polyadenylation signal (HDR/SV40pA) (sequences are listed in Supplemental Text). RT mixes were prepared using the superscript IV reverse transcriptase kit (Life Technologies, CA, USA) and PCR mixes using the Q5® High-Fidelity PCR Kit (New England Biolabs, MA, USA) following the manufacturer’s instructions. RT were performed in the following conditions: 25°C for 10◻min followed by 37◻°C for 50 min and 70°C 15◻min. PCR amplifications were performed in the following conditions: 98◻°C for 30◻sec followed by 35 cycles of 98◻°C for 10 s, 62◻°C for 30◻s, 72◻°C for 2◻min 30 s, with a 2 min final elongation at 72◻°C. PCR product sizes and quality were controlled by running gel electrophoresis and DNA fragments were purified using a QIAquick PCR Purification Kit (Qiagen, Hilden, Germany).

#### Cell transfection

HEK-293 cells were seeded into six-well cell culture plates one day prior to transfection. Cells were transfected with 2 μg of an equimolar mix of the five DNA fragments using lipofectamine 3000 (Life Technologies) following the manufacturer’s instructions. Each transfection was performed in five replicates. After incubating for 24 hrs, the cell supernatant medium was removed and replaced by fresh cell culture medium. Seven days post-transfection, cell supernatant medium was passaged two times using six-well cell culture plates of confluent C6/36 cells. Cells were subsequently inoculated with 100 μL of diluted (1/3) cell supernatant media, incubated 1 hr, washed with PBS 1X, and incubated 7 days with 3 mL of medium. Remaining cell supernatant medium was stored at −80◻°C. The second passage was used to produce virus stock solutions of DENV-1 1806 WT and mutant viruses. Transmission efficiency was assessed 21 days after an infectious blood meal containing the Parental, the Parental construct, the P10 strain, the P10 constructs (1 and 2) provided separately at a titer of 10^7^ FFU/mL.

### Statistical analyses

Statistical analyses were conducted using the STATA software (StataCorp LP, Texas, and USA). P-values>0.05 were considered non-significant. If necessary, the significance level of each test was adjusted based on the number of tests run, according to the sequential method of Bonferroni (74).

## Acknowledgments

The authors thank Pascal Delaunay (Centre Hospitalier Universitaire Nice, France), Ashgar Talbalaghi (Vector Control, Italy), Roger Eritja (Universitat de Barcelona, Spain), Vincent Robert (IRD, Montpellier) for providing mosquito eggs. This study was supported by a grant FP7-HEALTH HEALTH.2011.2.3.3-2 (grant number: 282378) “Dengue research Framework for resisting epidemics in Europe” (DENFREE), the French Government’s Investissement d’Avenir program, Laboratoire d’Excellence “Integrative Biology of Emerging Infectious Diseases” (grant n°ANR-10-LABX-62-IBEID), the European Union’s Horizon 2020 research and innovation programme under ZIKAlliance grant agreement No 734548 and the Institut Pasteur. PY was supported by the Pasteur-Paris University (PPU) program. LL received support from Agence Nationale de la Recherche (grant ANR-17-ERC2-0016-01), the City of Paris Emergence(s) program in Biomedical Research, and the European Union’s Horizon 2020 research and innovation programme under ZikaPLAN grant agreement No 734584. The funders had no role in study design, data collection and analysis, decision to publish, or preparation of the manuscript.

## Author contributions

LL and ABF designed the research. RB, SL, HJ, GPG, LM, GP, and FA performed experiments. LM, GP, PSY, GG, and MV provided reagents and analytical tools. AS and XDL provided expertise and feedback. RB, SL, HJ, GPG, GPP, and ABF analyzed the data. GPG, GPP, LL, and ABF wrote the manuscript. LL and ABF secure funding.

---

**Table S1.** Comparisons of infection rates and dissemination efficiencies between mosquitoes infected with two DENV-1 strains (1806 and 30A) and examined at different days post-infection (14 and 21).

| Mosquito population | Day post-infection | Fisher's exact test |  |
| --- | --- | --- | --- |
|  |  | Infection rate | Dissemination efficiency |
| Alessandria | 14 | 0.093 | 0.905 |
|  | 21 | 0.556 | 0.479 |
| Genoa | 14 | 0.077 | 0.022 |
|  | 21 | 0.035 | 0.376 |
| Cornella | 14 | 0.979 | 0.168 |
|  | 21 | 0.808 | 0.715 |
| Martorell | 14 | 0.496 | 0.001 |
|  | 21 | 0.006 | 0.002 |
| Nice | 14 | 0.191 | 0.929 |
|  | 21 | 0.371 | 0.463 |
| Saint-Raphael | 14 | 0.18 | 0.114 |
|  | 21 | 0.334 | 0.334 |

**Table S2.** Comparisons of infection rates and dissemination efficiencies between the 6 mosquito populations infected with a DENV-1 strain (1806 or 30A) and examined at a given day post-infection (14 or 21).

| DENV-1 | Day post-infection | Fisher's exact test |  |
| --- | --- | --- | --- |
|  |  | Infection rate | Dissemination efficiency |
| 1806 | 14 | 0.0001 | 0.0001 |
|  | 21 | 0.001 | 0.0001 |
| 30A | 14 | 0.0001 | 0.0001 |
|  | 21 | 0.0001 | 0.0001 |

**Table S3.** Comparisons of viral loads in mosquito heads between mosquitoes infected with two DENV-1 strains (1806 and 30A) and examined at different days post-infection (14 and 21).

| Mosquito population | Day post-infection | Wilcoxon Rank-Sum test |
| --- | --- | --- |
| Alessandria | 14 | 0.292 |
|  | 21 | 0.746 |
| Genoa | 14 | 0.533 |
|  | 21 | 0.836 |
| Cornella | 14 | 0.439 |
|  | 21 | 0.018 |
| Martorell | 14 | 0.073 |
|  | 21 | 0.304 |
| Nice | 14 | 0.183 |
|  | 21 | 0.857 |
| Saint-Raphael | 14 | 0.032 |
|  | 21 | 0.169 |

**Table S4.**
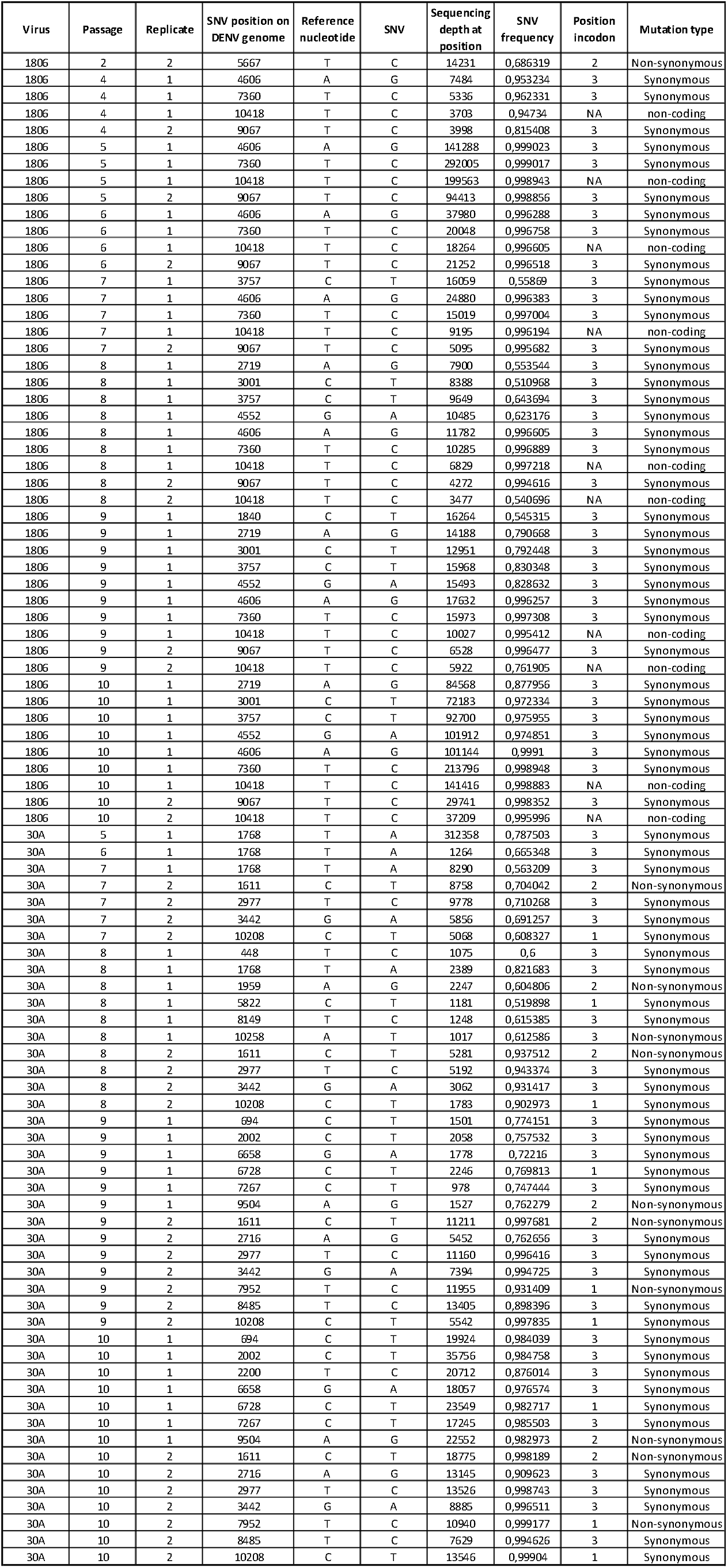
Position and estimated frequency of SNVs reaching consensus level detected in the mosquito-passaged samples.

**Table S5.**
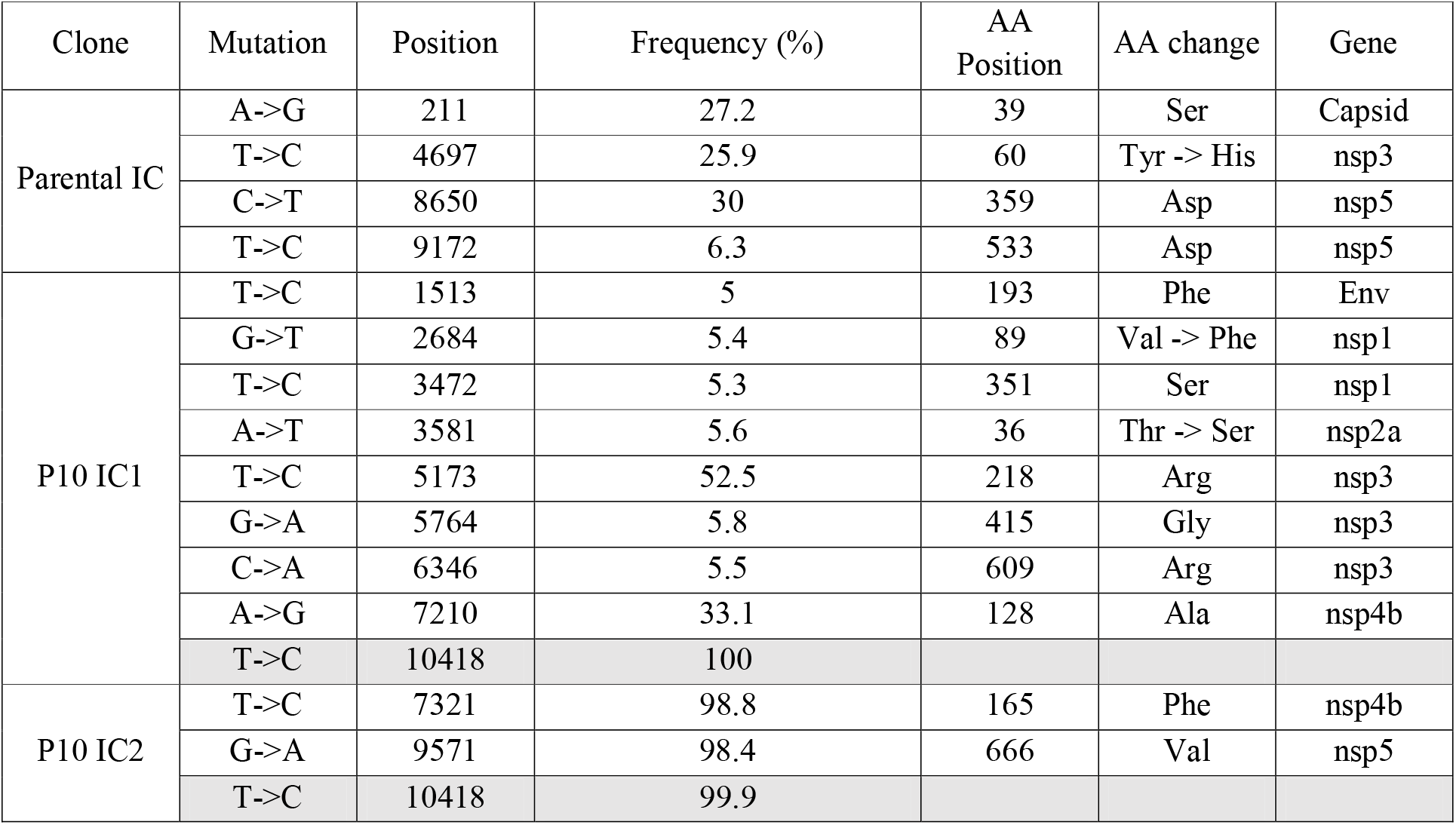
Sequences of the three reverse genetic constructs (Parental, P10 construct 1, P10 struct 2) using the deep sequencing method described in materials and methods. Only substitutions with a mutation frequency > 5% were considered significant for further analysis. The parental strain P0 was used as the reference sequence. The two P10 constructs presented the tation 10,418 (in grey) at a frequency close to 100% beside other mutations.

**Table S6.**
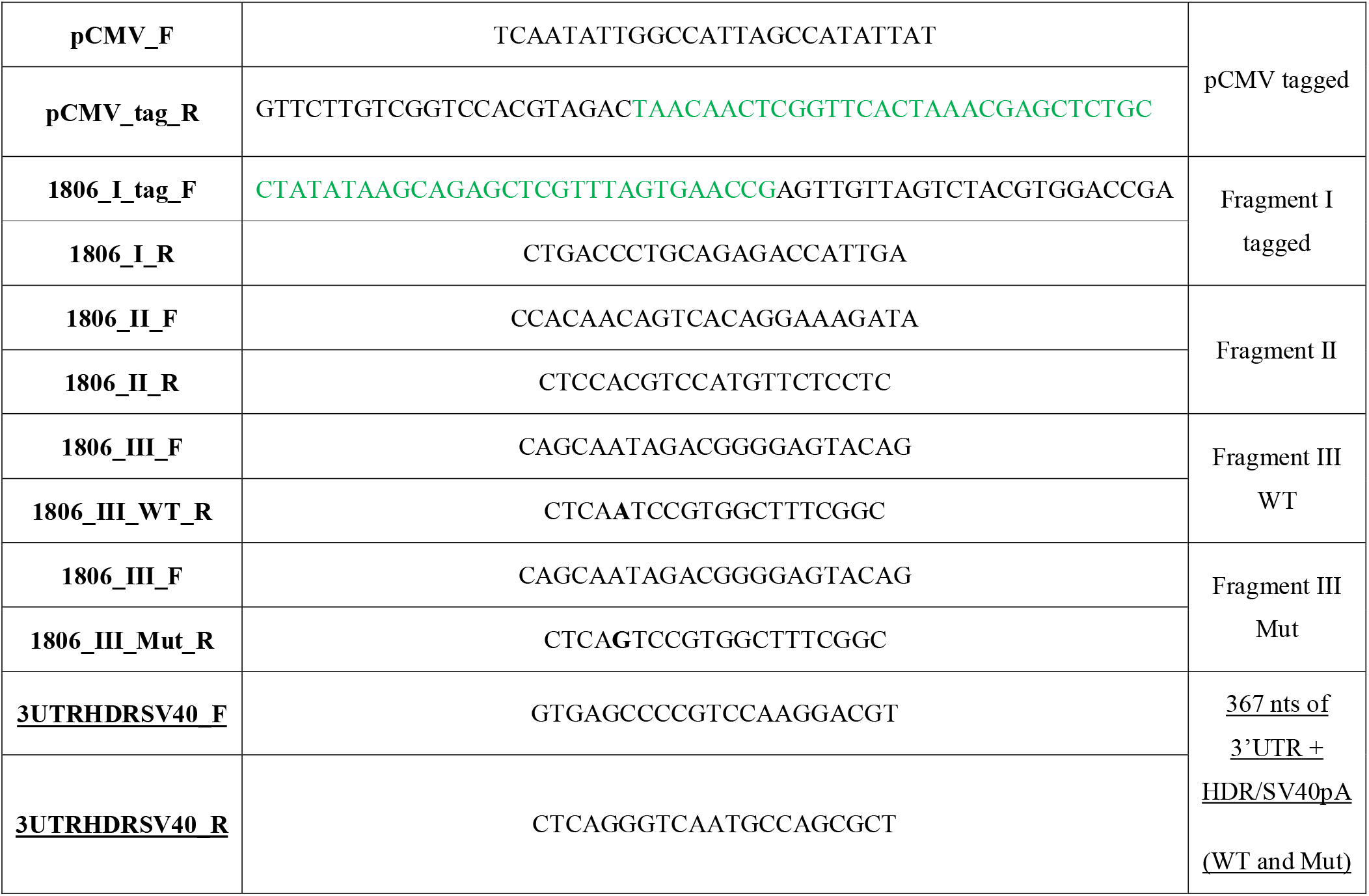
Primers used to generate subgenomic DNA fragments (ISA procedure)

**Figure S1.**
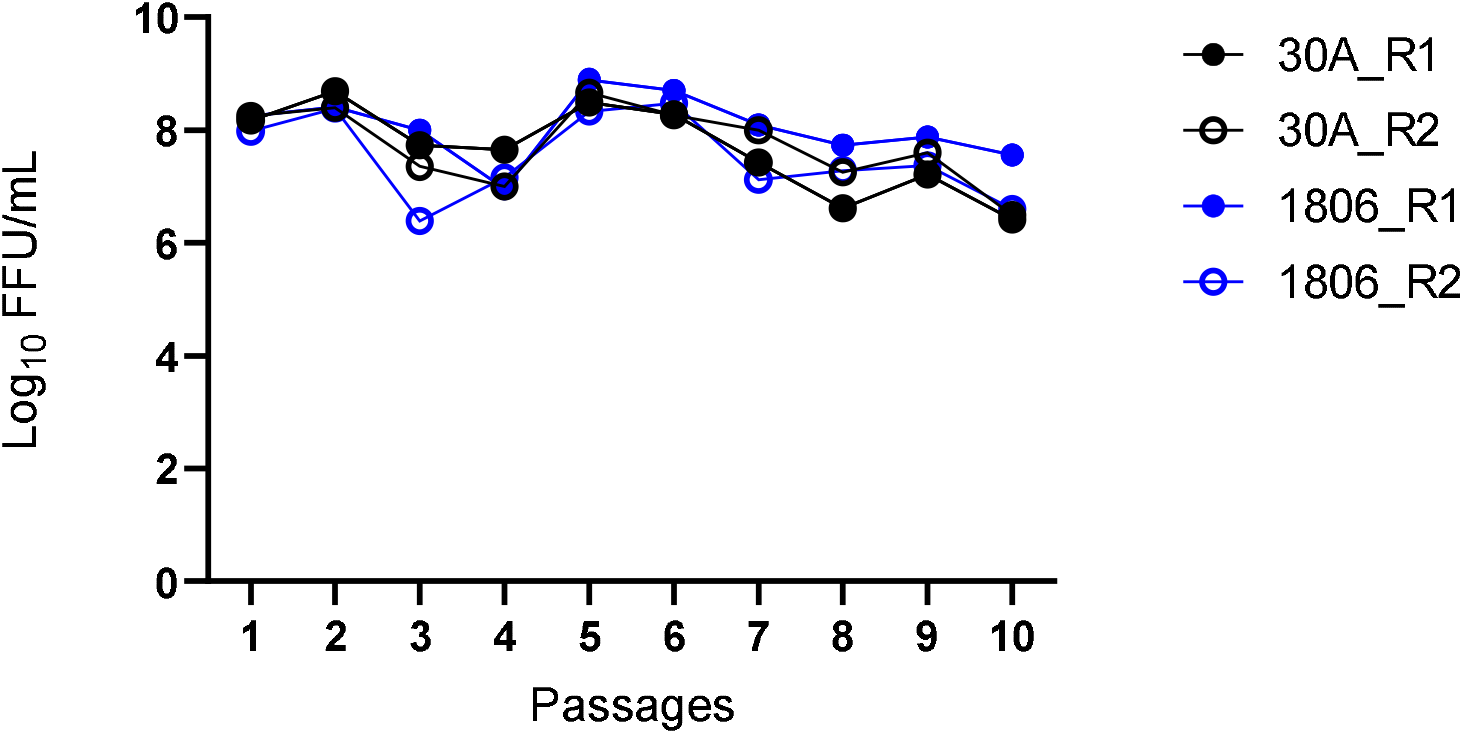
Viral titers of cell culture supernatants used to run passages in the experimental selection for DENV-1 adaptation to *Ae. albopictus*. Saliva were collected from 15-25 mosquitoes 19-21 days after infection and pooled to inoculate a monolayer of C6/36 *Ae. albopictus* cells. After 8 days at 28°C, cell culture supernatants were collected and provided to mosquitoes to run the next passage. Ten passages were performed. The supernatants were titrated by focus fluorescent assay on *Ae. albopictus* C6/36 cells. Viral titer was expressed in FFU/mL. Two biological replicates were performed for each viral strain, DENV-1 30A and DENV-1 1806.

**Figure S2.**
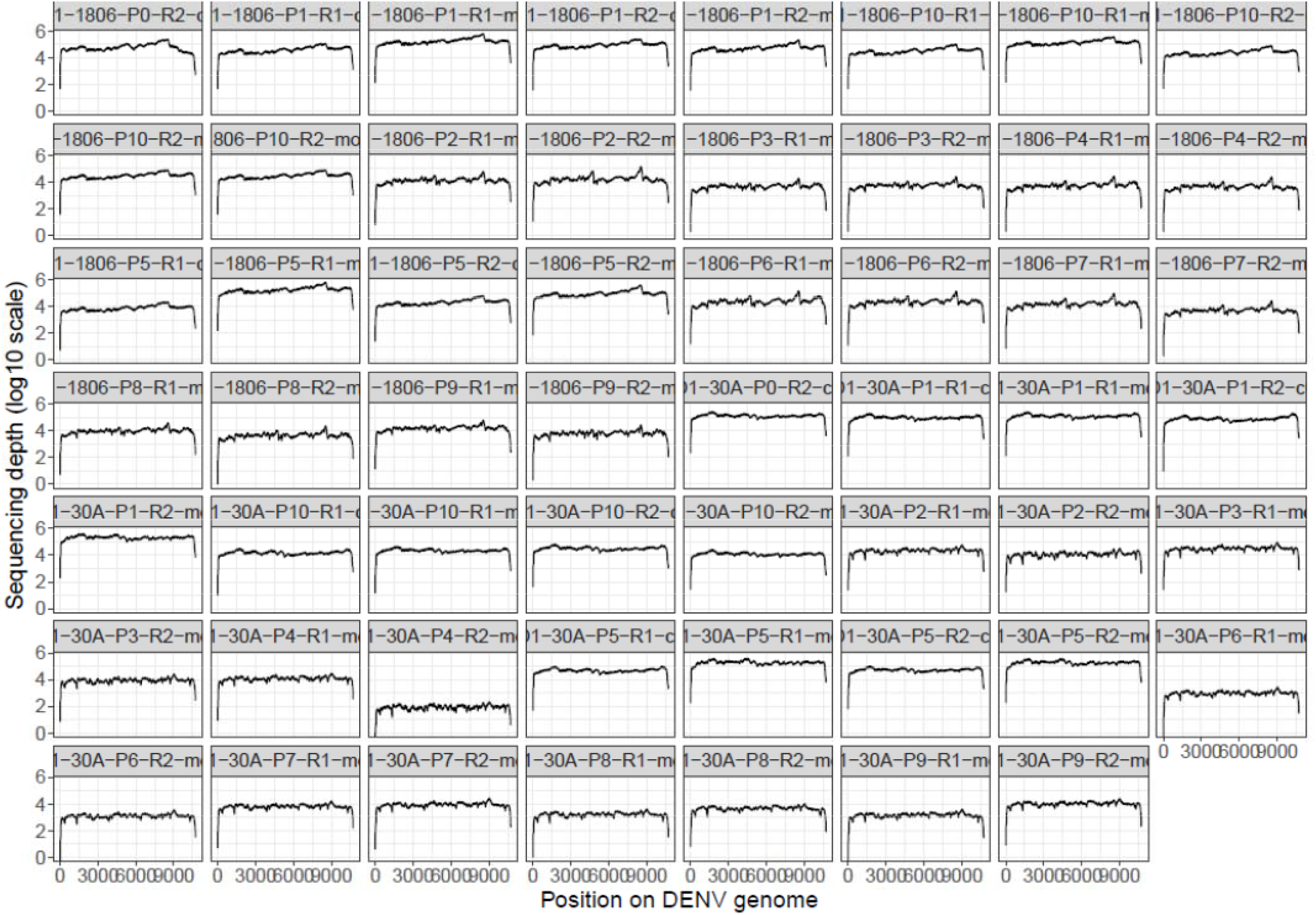
Sequencing coverage and depth by sample.

## Supplemental Text

Sequences of the *de novo* subgenomic DNA fragments used during the ISA procedure

### pCMV

GAATAAGGGCGACACGGAAATGTCACCCAACTGATCTTCAGCATCTTCAATATTGGCCATTAGCCAT ATTATTCATTGGTTATATAGCATAAATCAATATTGGCTATTGGCCATTGCATACGTTGTATCTATATCA TAATATGTACATTTATATTGGCTCATGTCCAATATGACCGCCATGTTGGCATTGATTATTGACTAGTTA TTAATAGTAATCAATTACGGGGTCATTAGTTCATAGCCCATATATGGAGTTCCGCGTTACATAACTTA CGGTAAATGGCCCGCCTGGCTGACCGCCCAACGACCCCCGCCCATTGACGTCAATAATGACGTATGTT CCCATAGTAACGCCAATAGGGACTTTCCATTGACGTCAATGGGTGGAGTATTTACGGTAAACTGCCCA CTTGGCAGTACATCAAGTGTATCATATGCCAAGTCCGCCCCCTATTGACGTCAATGACGGTAAATGGC CCGCCTGGCATTATGCCCAGTACATGACCTTACGGGACTTTCCTACTTGGCAGTACATCTACGTATTA GTCATCGCTATTACCATGGTGATGCGGTTTTGGCAGTACACCAATGGGCGTGGATAGCGGTTTGACTC ACGGGGATTTCCAAGTCTCCACCCCATTGACGTCAATGGGAGTTTGTTTTGGCACCAAAATCAACGGG ACTTTCCAAAATGTCGTAATAACCCCGCCCCGTTGACGCAAATGGGCGGTAGGCGTGTACGGTGGGA GGTCTATATAAGCAGAGCTCGTTTAGTGAACCG

### 367 last nucleotides of 3’UTR + HDR/SV40pA (WT; T)

GTGAGCCCCGTCCAAGGACGTAAAATGAAGTCAGGCCGAAAGCCACGGA**T**TGAGCAAGCCGTGCTG CCTGTGGCTCCATCGTGGGGATGTAAAAACCCGGGAGGCTGCAACCCATGGAAGCTGTACGCATGGG GTAGCAGACTAGTGGTTAGAGGAGACCCCTCCCTAGACATAACGCAGCAGCGGGGCCCAACACCAG GGGAAGCTGTACCTTGGTGGTAAGGACTAGAGGTTAGAGGAGACCCCCCGCACAACAACAAACAGC ATATTGACGCTGGGAGAGACCAGAGATCCTGCTGTCTCTACAGCATCATTCCAGGCACAGAACGCCA GAAAATGGAATGGTGCTGTTGAATCAACAGGTTCTGGCCGGCATGGTCCCAGCCTCCTCGCTGGCGC CGGCTGGGCAACATTCCGAGGGGACCGTCCCCTCGGTAATGGCGAATGGGACTCGCGACAGACATGA TAAGATACATTGATGAGTTTGGACAAACCACAACTAGAATGCAGTGAAAAAAATGCTTTATTTGTGA AATTAAGCGCTGGCATTGACCCTGAGGTTTACCCTCACAACGTTCCAGT

### 367 last nucleotides of 3’UTR + HDR/SV40pA (Mutant; C)

GTGAGCCCCGTCCAAGGACGTAAAATGAAGTCAGGCCGAAAGCCACGGA**C**TGAGCAAGCCGTGCTG CCTGTGGCTCCATCGTGGGGATGTAAAAACCCGGGAGGCTGCAACCCATGGAAGCTGTACGCATGGG GTAGCAGACTAGTGGTTAGAGGAGACCCCTCCCTAGACATAACGCAGCAGCGGGGCCCAACACCAG GGGAAGCTGTACCTTGGTGGTAAGGACTAGAGGTTAGAGGAGACCCCCCGCACAACAACAAACAGC ATATTGACGCTGGGAGAGACCAGAGATCCTGCTGTCTCTACAGCATCATTCCAGGCACAGAACGCCA GAAAATGGAATGGTGCTGTTGAATCAACAGGTTCTGGCCGGCATGGTCCCAGCCTCCTCGCTGGCGC CGGCTGGGCAACATTCCGAGGGGACCGTCCCCTCGGTAATGGCGAATGGGACTCGCGACAGACATGA TAAGATACATTGATGAGTTTGGACAAACCACAACTAGAATGCAGTGAAAAAAATGCTTTATTTGTGA AATTAAGCGCTGGCATTGACCCTGAGGTTTACCCTCACAACGTTCCAGT

